# A comparison of honeybee and bumblebee performance in matching-to-sample tasks

**DOI:** 10.64898/2026.01.31.703063

**Authors:** Romain Willemet

## Abstract

Reports of honeybees demonstrating abstract concepts like sameness and difference marked a pivotal development in comparative psychology. Subsequent studies expanded the scope of concept learning in honeybee cognition, yet most evidence relies on a single method: the delayed-matching-to-sample task using a Y-maze. Whether this setup is uniquely effective or if alternative approaches could yield similar results remains unresolved. Additionally, the failure of bumblebees (Bombus spp.) to complete this task, despite honeybees demonstrating success, remains unexplained. This study compared the performance of honeybees (*Apis mellifera*) and bumblebees (*Bombus terrestris*) across matching-to-sample tasks with varying degrees of physical continuity between sample and target stimuli. The objectives were twofold: to evaluate an alternative method for assessing concept learning in both species and to investigate potential species differences in such tasks. Contrary to prior findings, neither species succeeded at the reported proficiency levels in simultaneous matching-to-sample tasks. Moreover, bumblebees outperformed honeybees in one task. These results are consistent with an explanation based on species-specific differences in visual attention mechanisms, and underscore the need for further research on concept learning in social bees.

## Introduction

Abstract concept formation—the ability to generalize beyond the immediate perceptual properties of objects—has been considered a cornerstone of human cognition since the foundational work of early modern psychology (James et al., 1890). While methodological challenges, such as the influence of training-set size, have complicated studies in non-human animals (Katz & Wright, 2021), accumulating evidence indicates that concept learning occurs in multiple bird and mammal species (Zentall et al., 2008). This research has since expanded to insects, with evidence that honeybees (*Apis mellifera*), trained in a Delayed-Match-to-Sample (DMTS) paradigm, could generalize the concept of “sameness” to novel stimuli (Giurfa et al., 2001). The finding was pivotal for comparative psychology, as it revealed that a brain with fewer than one million neurons—a scale orders of magnitude smaller than those of species having successfully showed a capacity of concept learning until then (Herculano-Houzel et al., 2007, 2011; Olkowicz et al., 2016)—could support such cognitive capacity. Although computational models have since replicated aspects of these tasks with minimal neural architecture (Cope et al., 2018), the capacity for concept-learning in honeybees reveals a cognitive complexity that exceeds earlier expectations.

In the seminal paper by Giurfa et al. (2001), individual honeybees could enter a Y-maze by flying through a hole at the centre of a sample stimulus, and then choose between two target stimuli, only one of which was rewarded. The rewarded target stimulus was similar to the sample stimulus, while the unrewarded target differed from the sample stimulus. Giurfa et al. (2001) conducted a number of variants of this delayed-matching-to-sample (DMTS) task (Blough, 1959; Lind et al., 2015), using both visual (colour and pattern stimuli) and olfactory stimuli. They also tested honeybees in similar conditions, but with the rewarded target stimulus being the one that differed from the sample stimulus (i.e. a delayed-non-matching-to-sample, or DNMTS, task). The two major findings of these experiments are that (i) honeybees could improve their performance (measured as the proportion of correct choice) over several dozen trials and (ii) the bees maintained a better-than-chance level of performance during non-rewarded transfer tasks, in which new stimuli were used. This suggests that honeybees possess the capacity for concept learning. Remarkably, bees could do so even when the modality (visual or olfactory) of these new stimuli differed from that used during training. Using mazes of different lengths, Zhang et al. (2005) further reported that bees could retain information from the sample stimuli for up to five seconds, and even differentiate between multiple sample stimuli. The original Y-maze design has now been used to examine different cognitive abilities in honeybees, including the use of other concepts such as above and below (Avarguès-Weber et al., 2011), the simultaneous acquisition of two concepts (Avarguès-Weber et al., 2012), and numerical abilities (Howard et al., 2019).

A notable disparity exists between honeybees and bumblebees in performance on concept-learning tasks. Honeybees demonstrate proficiency in DMTS tasks including transfer tests, and have been shown to resolve complex mazes using sequential cues (Menzel, 2009; Zhang et al., 1996). Although bumblebees have rarely been tested on tasks requiring them to hold a cuing stimulus in memory, attempts have generally reported failures. Indeed, a replication of the original honeybee DMTS task (Giurfa et al., 2001) in bumblebees yielded chance-level performance after 80 trials (Sherry & Strang, 2015). Dale et al. (2005) found that bumblebees failed to solve a modified DMTS task using color cues alone, achieving above-chance performance only when a spatial component was added. While bumblebees showed limited success in a DMTS task using floral images (Thompson & Plowright, 2016), only a subset performed above chance, and spatial stability of rewarded targets may have aided learning. Despite documented differences in cognitive abilities, such as serial reversal learning (Sherry & Strang, 2015), the specific factors underlying this interspecies performance gap in DMTS tasks remain unresolved.

To examine this question, the present study evaluated bee performance in matching-to-sample tasks to investigate rule-based learning and elucidate species-specific differences between bumblebees and honeybees. Matching-to-sample tasks are conceptually analogous to delayed-matching-to-sample tasks, with the key distinction that the sample stimulus is present during choice. While the presence of the sample stimulus introduces alternative interpretative challenges (e.g. perceptual interferences), this reduces the memory load and theoretically lowers cognitive demand (Zentall & Smith, 2016). This reduction in task complexity provides an initial, simplified framework for evaluating species differences (Gentner et al., 2021).

In the experiments reported here, spatial continuity between sample and target stimuli was manipulated by varying the physical gap between them in a two-dimensional arrangement. Bees matched the sample to the target across three protocols. First, in the *full sample stimulus* experiment, target stimuli were integrated into the sample stimulus, with only the unrewarded target visually distinct. This simplified task was tested exclusively in bumblebees due to prior reports of poor performance in similar paradigms. Second, in the *surround sample stimulus* experiment, the sample stimulus enclosed the target stimuli, ensuring simultaneous access to both from all approach angles. Third, in the *central sample stimulus* experiment, the sample stimulus was positioned between the two target stimuli, matching the dimensions used in the original DMTS study by Giurfa et al. (2001). The latter two protocols were conducted in both bumblebees and honeybees. The two species were then compared in terms of the evolution of correct choice proportions and the strategies employed during decision-making.

## Material and methods

### Overview

The experiments were conducted at Royal Holloway, University of London, in England, between April-May 2020 and August 2022. Bee behavior was recorded during Matching-to-Sample experiments where the physical separation between the sample stimulus (indicating the rewarded color for that trial) and the two target stimuli was systematically varied to incrementally increase task complexity. Honeybees were tested in two experimental variants, and bumblebees in three. Individual bees, marked with acrylic paint on the thorax and upper abdomen for identification, completed 48 consecutive trials. Only the target stimulus matching the sample contained a rewarding sugar solution, while the non-matching target contained an aversive quinine solution. A choice was recorded as correct if the bee initially landed on the target matching the sample stimulus, and incorrect otherwise.

Transfer tests, typically involving unrewarded novel stimuli, are conventionally used to assess whether subjects generalize concepts beyond the training set, thereby confirming that performance improvements reflect conceptual understanding rather than mere familiarity with specific stimulus configurations (Giurfa, 2021; Wright et al., 2003). However, the introduction of new stimuli with altered salient features necessitates additional familiarization procedures (Giurfa et al., 2001), which can complicate the analysis of species-specific performance differences. This is because such procedures may introduce confounding variables, including differences in exploratory behavior and neophobia (see also Collado et al., 2021), potentially obscuring species-specific differences and increasing intra-species variability (Muller et al., 2010). Direct comparisons between species, such as those conducted here, are rare, and the absence of reported differences between the two species studies here likely reflects this gap. Furthermore, the use of transfer stimuli is contingent on color selection, ideally requiring multiple combinations of training and transfer stimuli to be tested (Banschbach, 1994; Chow et al., 2022). Given the primary aim of this study—to compare bumblebee and honeybee performance in matching-to-sample tasks—transfer tests were excluded. Pilot data indicated low performance in both species even after extensive training, rendering such tests unwarranted. By restricting stimuli to two consistent colors, the current experiments enabled direct performance comparisons between species under identical conditions.

Comparative studies require consideration of the factors that facilitate or constrain learning. Addressing this aspect allows for the development of species-specific, ecologically valid tests of concept learning. This paper centers on this aim, and none of the claims presented depend on the inclusion of transfer tests.

### Animals

#### Bumblebees

Four bumblebee colonies (*Bombus terrestris audax,* Biobest, Belgium) were used. Upon arrival, each colony was transferred in a plastic box (27x15x10cm) and placed in the experimental room following a 12h:12h light:dark photoperiod using daylight fluorescent tubes and high-frequency ballasts. Bees were provided daily with pollen deposited inside the nesting area.

#### Honeybees

The honeybees came from four observation hives situated in the campus the university. A nucleus colony of honeybees had been trained to feed on a feeder situated a few meters away from their hive and containing a 20% weight/weight sugar solution. On each experimental day, a single bee was transferred from the communal feeder towards the entrance of the outdoor testing arena using a cotton tip soaked with a 45% weight/weight sugar solution. This bee was then trained with the experimental protocol described below.

### Apparatus

The flight arena was made of plastic mesh and transparent acrylic glass walls (length, width: 55cm; height: 60cm) and covered by a plastic mesh ceiling. A magnetic black board (height: 27cm, width, 37cm) was fixed to a wall opposite the entrance hole, allowing us to display both the stimuli and the feeding stations (Figure 1.a). When bumblebees were tested, the arena was connected to the nest box with a GAWA type individual bee selector (Willemet, 2025). Honeybees entered the arena on their own by flying through the entrance hole. In both cases, the entrance hole was situated at the top of the arena so that the stimuli would appear in the ventral part of their visual field, the part most efficient for visual discrimination tasks (Lehrer, 1999; Morawetz et al., 2015; Wehner, 1972).

**Figure 1.**
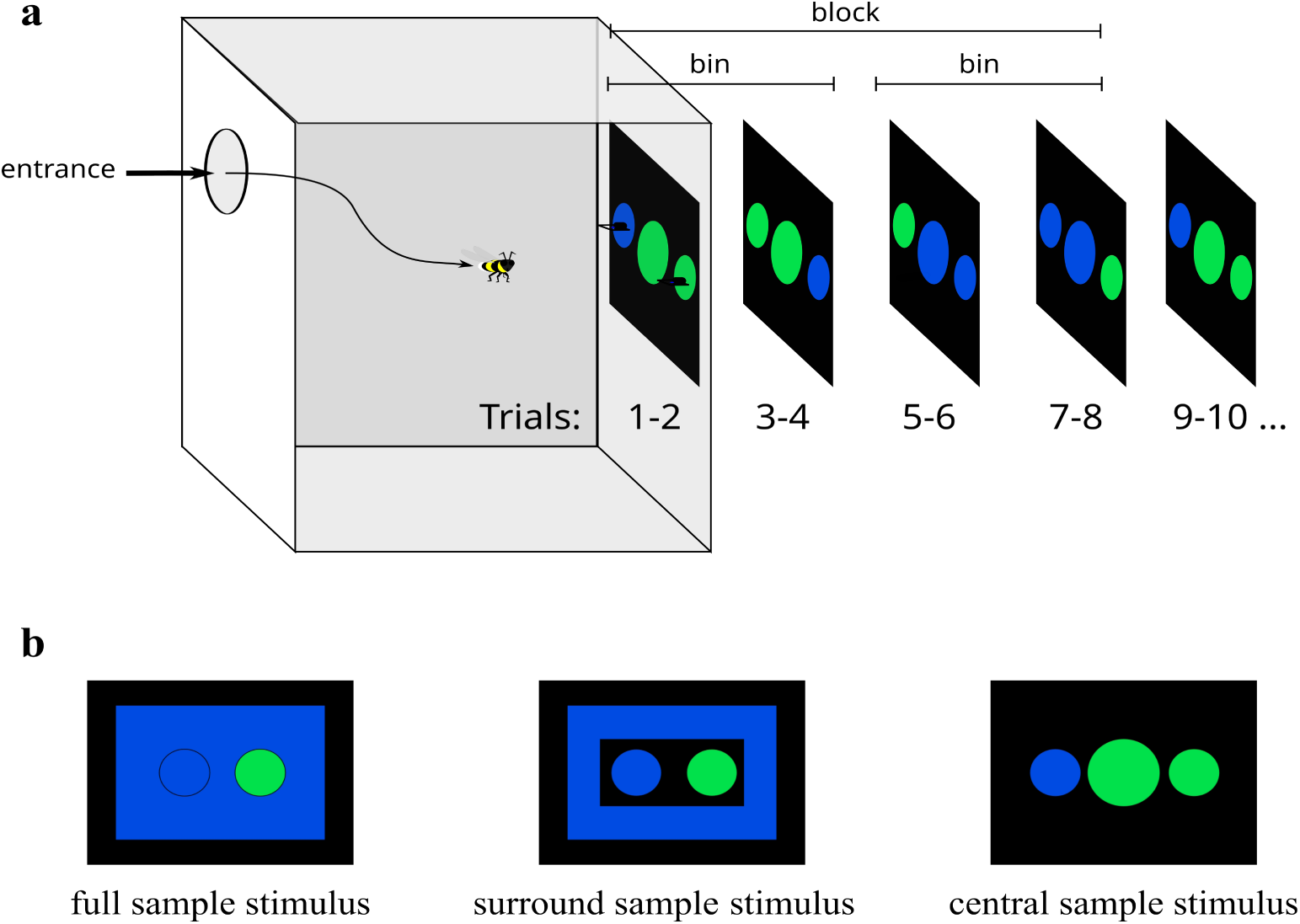
a. Representation of the arena and arrangement of the stimuli. Each combination (colour and side) of sample and target stimuli is presented twice in a row. That is, the colour of the sample stimulus changes every four trials and the side of the rewarding target stimulus changes every two trials. b. The three kinds of sample/target stimuli used in the experiments.

### Stimuli

The target stimuli were discs 7 cm in diameter. The two colours used for the stimuli were blue (RGB: 0,75,225) and green (RGB 0,225,75), printed on white paper and glued to flexible magnetic sheets. These colours were chosen because they had been shown to be well differentiated by bumblebees (Perry et al., 2016). Three shapes of blue and green sample stimuli were used in order to create a gradient of physical continuity between the sample and target stimuli (Figure 1.b):

1. The full sample stimuli were 29cm * 20.5cm rectangles. The target stimuli were positioned directly on them, so that there was a physical continuity between the sample and target stimuli, and only the incorrect stimulus (of a different colour) would stand out. Only the bumblebees were tested on this set-up due to the relative paucity, compared to honeybees, of reports of conditional foraging.
2. The surround sample stimuli resembled the full sample stimuli but included a central rectangular hole (19 cm × 10 cm) that allowed the target stimuli to be positioned on a dark background, introducing a physical discontinuity between the sample and target stimuli.
3. Finally, the central sample stimuli were discs 10 cm in diameter, placed between the target stimuli. The central sample condition thus had the smallest sample stimuli, over the most restricted area of the presentation panel, of the three conditions.

### Reward and punishment

The solutions (45° Bx or saturated quinine solution for reward and punishment, respectively) were deposited as drops (∼150 µl; i.e. sufficient to fill to saturation in a single visit) on a glass slide held in place by a small wooden support onto which the bees could land and access the solution. The aversive solution of quinine was preferred over simply not rewarding the alternative stimulus, as aversive reinforcement has been shown to improve performance in discrimination tasks in bees (Avarguès-Weber et al., 2010; Chittka et al., 2003). Both the microscope slide and the wooden platform were used only once per experiment, which precludes any odour cues from scent marks (Giurfa & Núñez, 1992) from influencing the results.

### Procedure

#### Pretraining

Before being tested, the bees were habituated to feed on a single stimulus of blue-green colour (RGB: 0,150,150). Bumblebees were trained in groups to forage on feeders resembling those used during the experiment with an unlimited access to a 30° Bx sugar solution. Honeybees were similarly trained, albeit individually. Only motivated bees, which, at the beginning at least, consistently took no more than approximately five minutes between foraging bouts, were selected for the experiments.

#### Training

Each experiment consisted of 48 trials (one trial starts with the bee entering the arena, and ends when it goes back to the nest/hive after it has ingested an unrestricted quantity of the sugar solution) and lasted from four to seven hours per subject. As in Giurfa et al. (2001), the sample stimulus was changed every four trials, and the rewarding side was changed every two trials. A bin corresponds to four consecutive trials with the same sample colour, and a block as two consecutive bins. Thus, a block comprised one bin for each colour, thereby controlling for potential colour preference biases. The experiments were recorded using a Logitech C525 webcam.

### Response variables and statistical analyses

A choice was recorded when the bee landed on the platform of a target stimulus. It was considered as correct when the bee landed on the target stimulus similar to the sample stimulus, and incorrect otherwise. Mixed models were computed to examine potential changes of performance over time, and chance level contrasts aimed to assess preferences or biases. More specifically, first to examine the bees’ overall performance (proportion of correct choices) over the course of the task, a binomial linear mixed effects model with block as predictor was computed with a logistic link function and subjects as random factor. Second, to examine whether the choice could be explained by a win-stay, lose-shift strategy (i.e. the tendency to continue exploiting a previously rewarding option and leave it when no longer rewarding), the effect of the order of choice-within-bin (first choice for each trial for the 1^st^, 2^nd^, 3^rd^ and 4^th^ trials within each bin) was examined. A binomial linear mixed effects model with choice-within-bin and block (of two bins) as predictors was computed. To investigate whether the bee’s performance during each training block was different from chance level, the proportions of correct choices were compared against a probability of 0.5 using one-sample t-tests. Note that p-value “correction” methods were not applied here as they are generally conservative for studies with limited sample size (VanderWeele & Mathur, 2019) and there are no *a priori* argument as to which error type (I or II) should be privileged. Rather, each individual p-values given here should be interpreted within the context of the other p-values and alternative lines of evidences (Johnson, 1999; Wasserstein et al., 2019). All analyses were performed using R Statistical Software (v4.1.2; R Core Team 2021).

### Ethical note

No licences or approval are required for experimental work on bumblebees and honeybees in the UK. However, care was taken taken to avoid negative impact on individual bees and their colonies. When possible, the bumblebee colonies were reused from ecophysiological experiments carried out by adjacent laboratories within the university, and for which only a small number of workers had been removed from the colony. On each experimental day, individual honeybees that had been tested previously and that were coming back to the tunnel were captured and temporary placed in a container with access to sugar water before being released at the end of the testing period.

## Results

### Honeybees

1- Honeybees – surround sample stimulus (13 bees)

#### Overall performance

The analysis revealed no main effect of block (Figure 2.a; Coef(β) = 0.07, SE(β) = 0.18, z=0.37, p=0.713, 95% confidence intervals [CI] = -0.30:0.44), implying that performance did not improve significantly over the course of the training. In fact, on average bees performed below chance level for the entire experiment (p-values for the 1^st^, 4^th^ and 6^th^ blocks<0.031). <u>Win-stay, lose-shift strategy</u>: The analysis revealed no main effect of block, no main effect of order of choice within bin, nor significant block x order of choice interaction within bin (p-values>0.804), see Figure 3.a.

**Figure 2.**
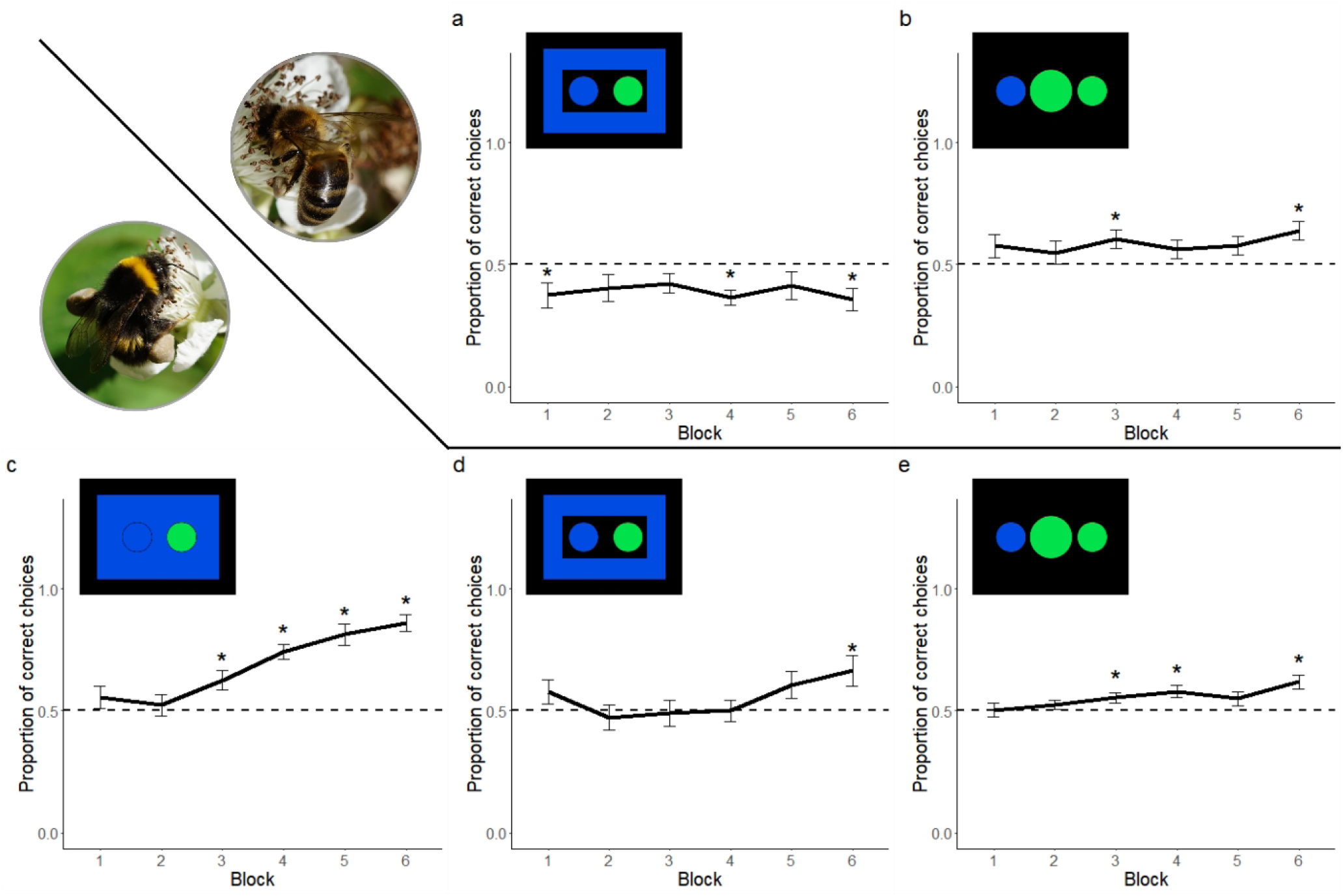
Proportion of correct choices across six blocks of eight trials for honeybees (a,b) and bumblebees (c,d,e). c: full sample stimulus, a,d: surround sample stimulus, b,d: central sample stimulus. Blocks for which the performance differs significantly from chance-level are indicated by a star.

**Figure 3.**
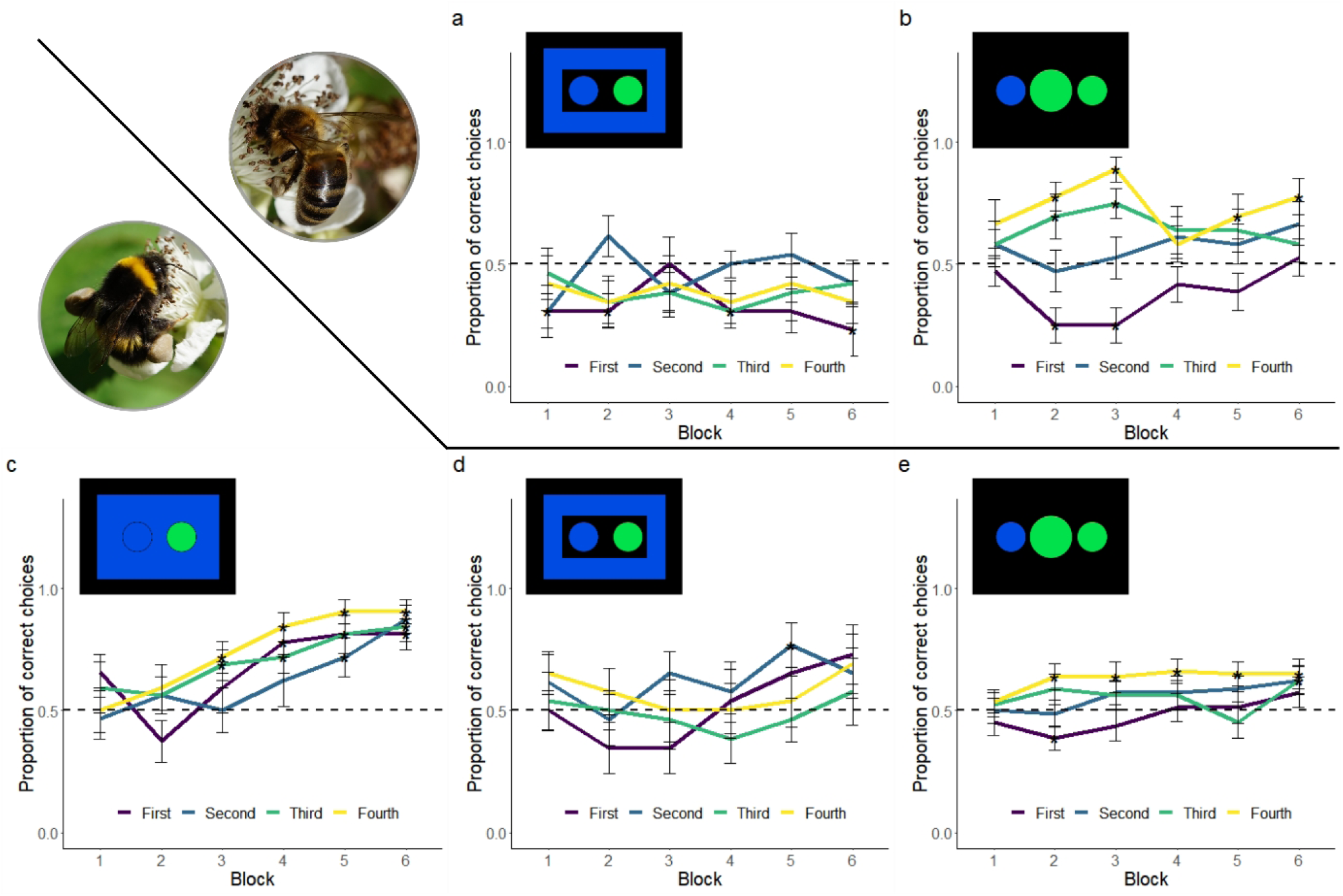
Proportion of correct choices within bins (first trial corresponds to a change in the sample stimulus, and third trial to a change in the side of the rewarding target stimulus, with second and fourth being identical to first and third trials, respectively) across six blocks, each containing two bins of four trials for honeybees (a,b) and bumblebees (c,d,e). c: full sample stimulus, a,d: surround sample stimulus, b,d: central sample stimulus. Blocks for which the performance differs significantly from chance-level for a given choice are indicated by a star. A lower than chance proportion of correct choices for the first and third trials within bins indicate that the bee tended to choose the side that was previously rewarding at the preceding trial, while a higher than chance proportion of correct choices for these trials suggest that bees respond to a change of the sample stimulus (first trial), or side of rewarding target stimulus (third trial).

2- Honeybees – central sample stimulus (10 bees)

#### Overall performance

Honeybees did not improve significantly over the course of the training (Figure 2.b, no main effect of block; p=0.090). <u>Win-stay, lose-shift strategy:</u> The analysis revealed no main effect of block, nor significant block x order of choice interaction within bin (p-values>0.119) but a significant effect of order of choice within bin (Coef(β) = 1.06, SE(β) = 0.25, z=4.29, p<0.001, 95% CI = 0.59:1.56). This suggests a win-stay, lose-shift strategy, visible in Figure 3.b.

### Bumblebees

1- Bumblebees – full sample stimulus (16 bees)

#### Overall performance

Bumblebees improved their performance over the course of the experiment. The analysis revealed a significant effect of block (Figure 2.c, Coef(β) = 0.65, SE(β) = 0.17, z=3.89, p<0.001, 95% CI = 0.34:1.00). <u>Win-stay, lose-shift strategy:</u> The analysis revealed no main effect of order of choice within bin, nor significant block x order of choice interaction within bin interaction (p-values>0.517) but a significant effect of block (Coef(β) = 0.34, SE(β) = 0.16, z=2.09, p=0.037, 95% CI = 0.03:0.66). This suggests an overall improvement of performance irrespective of the order of presentation within a bin, see Figure 3.c.

2- Bumblebees – surround sample stimulus (13 bees)

#### Overall performance

The analysis revealed no main effect of block (Figure 2.d, p=0.508). However bumblebees performed above chance level during the 6^th^ block (p=0.022). <u>Win-stay,</u> <u>lose-shift strategy</u>: The win-stay, lose-shift strategy analysis revealed no main effect of order of choice within bin, nor significant block by order of choice within bin interaction (p-values>0.062) but a main effect of block (Coef(β) = 0.52, SE(β) = 0.18, z=2.88, p=0.004, 95% CI = 0.17:0.89), see Figure 3.d.

3- Bumblebees – central sample stimulus (40 bees)

#### Overall performance

The analysis revealed a significant main effect of block (Figure 2.e, Coef(β) = 0.15, SE(β) = 0.08, z=1.96, p=0.050, 95% CI = 0.00:0.30). <u>Win-stay, lose-shift</u> <u>strategy</u>: The analysis also revealed a significant main effect of block (Coef(β) = 0.34, SE(β) = 0.11, z=3.14, p=0.002, 95% CI = 0.13:0.55). In addition, there was a significant main effect of order of choice within bin (Coef(β) = 0.44, SE(β) = 0.15, z=2.84, p=0.005, 95% CI = 0.14:0.74). There was no significant block by order of choice within bin interaction. Overall, this suggests that, like honeybees, bumblebees used a win-stay, lose-shift strategy (albeit in a less pronounced way, compare Figure 3.b and 3.e.).

## Discussion

This series of experiments yielded two primary findings: honeybees exhibited lower performance than previously reported, and species-specific differences emerged, with bumblebees outperforming honeybees. Results for each species are addressed separately below, followed by an integrated discussion.

### Honeybees

Honeybees have been reported to succeed in a variety of matching-to-sample tasks (Zhang et al., 2012). This ranges from oddity and non-oddity discrimination between sets of stimuli (Muszynski & Couvillon, 2015) to delayed-matching-to-sample (Brown et al., 1998; Couvillon et al., 2003; Giurfa et al., 2001; Zhang et al., 2005), and to symbolic delayed-matching-to-sample (Cooke et al., 2007; Howard et al., 2019; Moreno et al., 2012), including when the sensory modality of the sample stimulus differs from that of the target (Couvillon & Bitterman, 1988; Giurfa et al., 2001; Srinivasan et al., 1998) or when the sample stimulus gave information on the type of motor action to execute (Zhang et al., 1996).

The surround and central sample stimulus experiments presented here were variations of a simultaneous matching-to-sample task, with both sample and target stimuli continuously available. However, honeybees failed to achieve performance levels reported in previous studies. First-choice accuracy within blocks of four trials under the same sample stimulus remained at or below chance for both surround and central sample groups, indicating honeybees did not use the concept of sameness in their decision-making. In the surround sample stimulus experiment, performance remained below chance across blocks and within trial bins, suggesting bees responded to rewarding color and side without considering the sample stimulus.

Results differed in the central sample stimulus experiment, where performance improved, driven primarily by a color-based win-stay lose-shift strategy. Significant above-chance performance emerged only in the final block, likely due to responses to sample stimulus changes in the first trial of each bin. This could reflect either flexible attention to stimulus configurations or emerging recognition of the sample stimulus’s informational value. The former interpretation contrasts with multiple reversal learning studies, where honeybee performance declines after repeated reversals (Helversen, 1974; Menzel, 1968). The sample stimulus may have provided contextual cues that partially reset learning at each change. The latter interpretation is notable because performance improvement occurred only after more than 40 trials, exceeding the trial counts typically reported in DMTS experiments (e.g., Avarguès-Weber et al., 2011) but aligning with Giurfa et al. (2001). This underscores the need to investigate the effect of training set size on same-different concept learning (Katz & Wright, 2021).

Studies using trial-unique procedures with free-flying honeybees have reported rapid performance increases when rewards were associated with odd stimuli (Muszynski & Couvillon, 2015, 2020). However, direct reward and punishment availability in these studies raises questions about the influence of subtle sensory detection (Guiraud et al., 2018). Here, the lack of consistent improvement suggests that no such cue was used. Additionally, such cues would affect all trials equally, which contradicts the evidence of a win-stay, lose-shift strategy demonstrated here.

Of particular interest, performance in the central sample stimulus condition—most similar to Giurfa et al.’s (2001) original DMTS demonstration—did not surpass levels reported by Giurfa et al. (2001) after a comparable number of trials despite the fact that both the sample and target stimuli were presented simultaneously rather than with a time delay. The MTS design, although seemingly simpler than the DMTS design, appeared more challenging for honeybees.

### Bumblebees

The full sample stimulus experiment indicates that bumblebees rapidly learn either to select a target stimulus matching the sample stimulus color or to avoid the most visually distinct target in a two-color paradigm, though the present design does not distinguish between these possibilities. Increasing the spatial separation between sample and target stimuli in the surround sample experiment reduced bumblebee performance from over 80% correct choices at the end of training to approximately 65% (Figure 2.c.d). However, analysis of performance patterns within bins of four trials revealed a progressive improvement across all choices within individual bins, suggesting that bumblebees did not rely solely on previously rewarded side or color cues. This contrasts with the central sample stimulus experiment, where evidence of a weak color-based win-stay, lose-shift strategy was observed.

While the surround sample stimulus results suggest bumblebees may have learned the presentation rule (i.e., matching the sample stimulus), the absence of transfer stimuli precludes definitive conclusions. Bumblebees are known to integrate visual properties of complex stimuli (Fauria et al., 2000) and use color as a discriminative cue in operant conditioning (Mirwan et al., 2015). They also adjust decisions based on contextual information in dual-choice trials (Fauria et al., 2002), implying a theoretical capacity to learn the four sample-target combinations used here, rather than the relational rule between stimuli.

MaBouDi et al., (2020) reported that bumblebees improved performance in a task requiring attention to relative stimulus size, initially learning a relational rule before shifting to a win-stay, lose-shift strategy after 20 foraging bouts. While this may indicate concept learning, the rapid initial improvement followed by an abrupt cognitive shift—and the persistence of chance-level second choices—warrants further investigation.

Overall, these findings suggest that bumblebees may be able to improve performance in concept learning tasks under certain conditions, though further research is required to obtain conclusive evidence. The contrast between the surround sample stimulus experiment, which aligns with concept learning, and the central sample stimulus experiment, which reflects a win-stay lose-shift strategy, highlights the influence of experimental design on observed outcomes.

### General discussion

The experiments presented here demonstrate distinct performance levels between honeybees and bumblebees in matching-to-sample tasks. This corroborates earlier findings suggesting cognitive and behavioural difference between these two model species (Sherry & Strang, 2015). However, the observation that bumblebees outperformed honeybees under identical conditions contrasts with earlier findings, where honeybees succeeded and bumblebees struggled in delayed matching-to-sample (DMTS) tasks involving a temporal delay between sample and target stimuli.

The present results also contrast with several reports of concept learning in honeybees in matching-to-sample experiments. These previous experiments relied on procedures that differ from the set of experiment presented here, such as having a rewarded sample stimulus (Cooke et al., 2007; Couvillon et al., 2003), a trial-unique procedure, and with the stimuli presented horizontally (Muszynski & Couvillon, 2015). These procedures are known (Wolf et al., 2015), or expected from the work with other species (e.g. set size: (Overman Jr & Doty, 1980; Wright et al., 2018), to improve performance on learning tasks in bees. However, Giurfa et al. (2001) reported significant performance improvements in honeybees on a DMTS task despite not incorporating any of these procedural enhancements. This discrepancy suggests that the absence of a time delay between the presentation of the sample stimulus and of the target stimuli does not necessarily simplify the task for honeybees.

By contrast, bumblebees showed marked performance improvements in the full sample stimulus experiment and partial success in experiments with physical discontinuity between sample and target stimuli. This indicates that simultaneous presentation of stimuli may facilitate learning in bumblebees, who have shown no strong evidence of performance improvements in DMTS tasks so far (Sherry & Strang, 2015). However, the absence of transfer tests, chosen here to facilitate inter-species comparisons, precludes definitive conclusions about their capacity for abstract relational learning.

In all cases, the relatively low correct choice scores after 48 trials suggest that more extensive training may be necessary to fully assess the cognitive capacities of bumblebees. Notably, studies in birds and primates often involve hundreds or thousands of trials (Wright et al., 2017), whereas bee studies typically require only a few dozens. This discrepancy highlights the need for further research to determine the comparability of behavioral performance between insects and vertebrates.

While overall performance of bumblebees and honeybees in the central sample stimulus experiments appears comparable, the degree to which each species employs a win-stay, lose-shift strategy varies (Figure 3.b.e.). Future experiments should include detailed analyses of choice patterns to identify underlying cognitive strategies. Interestingly, both the evidence of colour-based win-stay, lose-shift strategy and the low performance in the honeybee surround sample suggest that bees failed to learn to switch sides every two trials, addressing a common criticism of this procedure. Larger sample sizes per group are necessary to ensure generalizability, given documented individual variability in concept learning tasks (Guiraud et al., 2018). Latency measurements before choice may also help by providing insight into cognitive processes (Chittka & Spaethe, 2007; MaBouDi et al., 2023; Willemet, 2026).

In addition, the use of blue and green stimuli, though well-associated in differential conditioning (Avarguès-Weber et al., 2010; MaBouDi, Marshall, et al., 2020; Mancini et al., 2018; Perry et al., 2016), may have influenced results. Pilot experiments indicated that naïve bumblebees quickly learn both colors in discrimination tasks, but color-specific effects on learning remain unclear (Avarguès-Weber & Giurfa, 2014; Kirkerud et al., 2017). Thus, generalizability requires testing multiple color combinations (Banschbach, 1994; Chow et al., 2022).

More generally, a deeper understanding of concept learning in honeybees and bumblebees necessitates further exploration on the factors influencing performance. Research in this area, such as the influence of set size, remains limited, even for honeybees, despite demonstrations that results obtained with the Y-maze protocol generally used in DMTS tasks can be subject to misinterpretations (Guiraud et al., 2018). Detailed response pattern analyses are essential to determine which species can resolve concept tasks and how they do so (Katz et al., 2002). For example, capuchin and rhesus monkeys both demonstrate abstract concept learning, but rhesus monkeys require additional procedural steps (touching the sample stimulus several times before the choice, Wright et al., 2003).

Beyond procedural requirements, the observed species differences may extend beyond methodological variations to reflect deeper cognitive disparities. Indeed, it is possible that the peculiar pattern of differences between honeybees and bumblebees in their capacity to resolve MTS and DMTS tasks is mainly explained by the difference between these two species in their capacity for parallel visual attention. The bee brain seems to have limited parallel visual information processing, which prevents bees to see “at a glance” (Nityananda et al., 2014; Skorupski et al., 2006). Instead, bees have to actively scan their environment for visual cues (MaBouDi et al., 2025). Interestingly, honeybees and bumblebees differ in that respect. Bumblebees seem able to perform parallel-like search on a visual task, while honeybees appear unable to do so (Morawetz & Spaethe, 2012). The honeybee’s serial visual attention mechanism may enhance performance in delayed matching-to-sample tasks but reduce effectiveness in matching-to-sample tasks, whereas the bumblebee’s parallel-like mechanism could produce the opposite outcome, potentially constrained by working memory limitations. This suggests that a seemingly limiting mechanism (lack of parallel visual attention) may allow honeybees to outperform bumblebees on a complex cognitive task (DMTS), despite the latter having a more sophisticated mechanism. This hypothesis is supported by the present observation that performance within species varied depending on how sample stimuli were presented. If true, such differences would be significant, as it would describe two closely related species that differ in their capacity to solve a kind of cognitive tasks due to having different cognitive limitations.

In all cases, the cognitive differences between honeybees and bumblebees represent a source of insight into how distinct visual attention mechanisms and working memory constraints shape task-specific performance. These differences also highlight the need for varied experimental designs to disentangle procedural influences from fundamental cognitive processes in concept learning tasks.

## Acknowledgements

The author is grateful to Ellouise Leadbeater for her input during the experimental phase and manuscript development, and to two anonymous reviewers for their comments on a previous version.

